# Visual acuity and egg spatial chromatic contrast predict egg rejection behavior of American robins

**DOI:** 10.1101/2020.05.21.109405

**Authors:** Alec B. Luro, Esteban Fernández-Juricic, Patrice Baumhardt, Mark E. Hauber

## Abstract

Color and spatial vision are critical for recognition and discrimination tasks affecting fitness, including finding food and mates and recognizing offspring. For example, as a counter defense to avoid the cost of raising the unrelated offspring of obligate interspecific avian brood parasites, many host species routinely view, recognize, and remove the foreign egg(s) from their nests. Recent research has shown that host species visually attend to both chromatic and spatial pattern features of eggs; yet how hosts simultaneously integrate these features together when recognizing eggs remains an open question. Here, we tested egg rejection responses of American robins (*Turdus migratorius*) using a range of 3D-printed model eggs covered with blue and yellow checkered patterns differing in relative square sizes. We predicted that robins would reject a model egg if they could visually resolve the blue and yellow squares as separate features or accept it if the squares blended together and appeared similar in color to the natural blue-green color of robin eggs as perceived by the avian visual system. As predicted, the probability of robins rejecting a model egg increased with greater sizes of its blue and yellow squares. Our results suggest that chromatic visual acuity and viewing distance have the potential to limit the ability of a bird to recognize a foreign egg in its nest, thus providing a limitation to host egg recognition that obligate interspecific avian brood parasites may exploit.

## Introduction

Animals use spatial chromatic cues when visually detecting predators and prey, choosing mates, and recognizing their own species and close kin (Caves et al., 2018; Cuthill et al., 2017). How animals perceive and discriminate between spatial chromatic visual cues when making decisions relevant to evolutionary fitness is a fundamental question of visual ecology (Cronin et al., 2014; Endler and Mappes, 2017). Behavioral experiments of the perception and discrimination of stimuli that vary both chromatically and spatially have been conducted in a variety of animal taxa, including honeybees *Apis mellifera* (Giurfa et al., 1996; Srinivasan and Lehrer, 1988), lizards (Fleishman et al., 2017), and birds (Lind and Kelber, 2011; Potier et al., 2018). In many of these studies, results from a limited number of captive animals subject to extensive training under controlled laboratory settings are considered to be representative and relevant to the model species’ natural behaviors and ecologies. Despite the potential for more direct inference of the effect of visual perception and discrimination on fitness in the wild, few studies have tested animals’ perception, discrimination, and behavioral responses to ecologically and evolutionarily relevant visual stimuli in their natural environments (Endler and Mappes, 2017).

Obligate avian brood parasitism is a rare breeding strategy (~1% of all bird species) in which brood parasites lay their eggs in the nests of different species. In response, potential hosts either recognize and reject the foreign egg(s) and/or nestling(s) or raise the parasitic offspring at a cost to their own fitness (Davies, 2000). Visually discriminating own vs. parasitic eggs can be an exceptionally challenging task when brood parasites produce highly mimetic eggs (Stoddard and Hauber, 2017). As a result, hosts may commit recognition errors and mistakenly reject their own eggs, rendering egg rejection a risky defense strategy with potentially severe consequences to host fitness (Davies et al., 1996; Lotem et al., 1995).

Arguably, avian brood parasitism research to test how birds recognize eggs is at the forefront avian visual perception and discrimination experiments conducted in the wild (Stoddard and Hauber, 2017). Model plaster (Davies and Brooke, 1989; Honza et al., 2007; Rothstein, 1982) or 3D printed eggs (Igic et al., 2015) can be systematically painted with ranges of natural avian colors (Canniff et al., 2018; Hauber et al., 2015) and spotting patterns (Dainson et al., 2017; Hanley et al., 2019; Luro et al., 2018) and placed into host species’ nests to test if the attending parents discriminate and recognize the model eggs by touching them (Soler et al., 2017), setting them aside within the nest, or removing them from the nest. Recent work combining visual modeling (Vorobyev and Osorio, 1998) with foreign-egg rejection experiments has demonstrated that 1) birds have a perceptual bias towards responding to natural blue-white-brown egg color gradients (Abolins-Abols et al., 2019; Manna et al., 2020), but do not respond predictably to purple-green color gradients not found among natural eggs (Hanley et al., 2017); 2) egg background color and egg spotting presence (and/or color) combine together as a multicomponent cue and can greatly increase or decrease egg rejection responses, depending on the host species’ own egg appearance (Dainson et al., 2017; Hanley et al., 2019; Luro et al., 2018); and 3) host species may use achromatic patterning and spatial features such as egg spotting and scrawling when recognizing and rejecting foreign eggs (Spottiswoode and Stevens, 2010; Stoddard et al., 2014). Egg rejection experiments require neither captivity nor extensive training and allow for individually repeated and/or population-wide tests of avian visual discrimination capabilities in an ecologically relevant context.

Despite great advances in our knowledge of how birds recognize foreign eggs within their nests using visual cues including egg size, shape, color and spotting, no study to date has examined whether egg spatial and chromatic features may be simultaneously integrated by the host’s visual system to influence egg recognition. For instance, how well can birds resolve details of egg spotting pattern and color when viewing eggs from various distances? Do some egg spotting patterns blend in with each other and with the egg background from the bird’s point of view? Here, we tested if and how American robins *Turdus migratorius* (hereafter: robins), a robust rejecter of parasitic brown-headed cowbird *Molothrus ater* eggs (Rothstein, 1982), respond to differences in spatial chromatic contrast when viewing and deciding to accept or reject a foreign egg. The underlying assumption is that visual acuity (i.e., visual spatial resolution) can limit the ability of birds to resolve details in the chromatic spatial pattern of eggs, as has been found in diverse taxa under different ecological conditions (Caves et al., 2018). We used avian visual modelling and visual acuity estimates for lateral and binocular vision of American robins to create model eggs with spatial chromatic patterns along a gradient of increasing difficulty for robins to visually resolve the patterned eggs as different from natural robin eggs. To this end, we i) designed a range of 3D printed model eggs and covered them with a suite of novel, blue and yellow checkered patterns, where both colors were present in equal proportions but differed in their relative inter-square distances to one another across model eggs, and ii) estimated a spatial chromatic discrimination threshold distance at which the blue and yellow squares should blend from the robin’s visual perspective and appear similar to the natural robin egg’s immaculate blue-green coloration. We predicted that robins would reject the model egg when they could resolve the blue and yellow squares as separate features, but would accept the artificial egg when they could not resolve the squares apart (i.e., if the blended color of blue and yellow squares appeared similar to the blue-green color of natural robin eggs).

## Methods

### Experimental Egg Pattern Design

We generated test checkerboard patterns of various combinations of blue and yellow colored squares, whose centers were spaced 0.15 mm apart from one another and printed the test color patterns on transferrable decal paper (Sunnyscopa Film-Free Waterslide Decal Paper Multi Use 8.5×10in [216×254mm], Sunnyscopa, South Korea) using an HP Color LaserJet Pro MFP M281fdw with HP202A black, yellow, cyan and magenta toners (HP Development Company, L.P. USA). Our printer had a color printing resolution of up to 600 × 600 dpi, and we printed patterns at 300 dpi resolution in tagged image file format (TIFF). We took spectral reflectance measurements of the printed test color patterns using an Ocean Optics JAZ spectrometer (Ocean Optics, Inc. USA) with a 400μm fiber optic cable and a reflectance probe fitted with a black rubber cap to block external light. We measured a 12.57 mm^2^ circular area of the pattern from a 10 mm distance at 90° angle. The reflectance probe had an acceptance angle of 24.8°, and therefore the reflectance measurements were of both blue and yellow colors together (using Eqn 1.0, 24.8° > 0.015° angle of single 0.150mm blue or yellow square measured from 10mm away).

*Eqn 1.0* (Swearer, 2011)

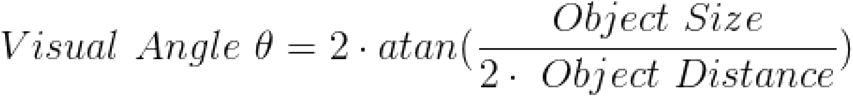

We used a receptor-noise limited model (Vorobyev and Osorio, 1998) to calculate avian-perceived chromatic and achromatic Just Noticeable Differences (JNDs) between our “blended” test blue and yellow pattern, the blue color alone, the yellow color alone, and a range of N = 22 natural robin eggs, using reflectance spectra data from Croston and Hauber (2015a) to determine the best printed blue/yellow combination for mimicking natural robin egg colors (Fig. 1). We used peak sensitivities of UVS, SWS, MWS and LWS cone photoreceptors, cut-off wavelengths (λ_cut_) for photoreceptors’ respective oil droplets, relative photoreceptor densities, and ocular media transmittance of the congeneric common (European) blackbird *Turdus merula* (Hart et al., 2000). The model assumed robins viewed the eggs under bright daylight conditions (D65 irradiance spectrum from Maia et al., 2019), eggs were viewed against the background of a typical dry brown grass lining in natural robin nests (nest reflectance data were sourced from Aidala et al., 2015), and a Weber fraction value of 0.1 for calculating photoreceptor noise values (Olsson et al., 2018).

**Figure 1.**
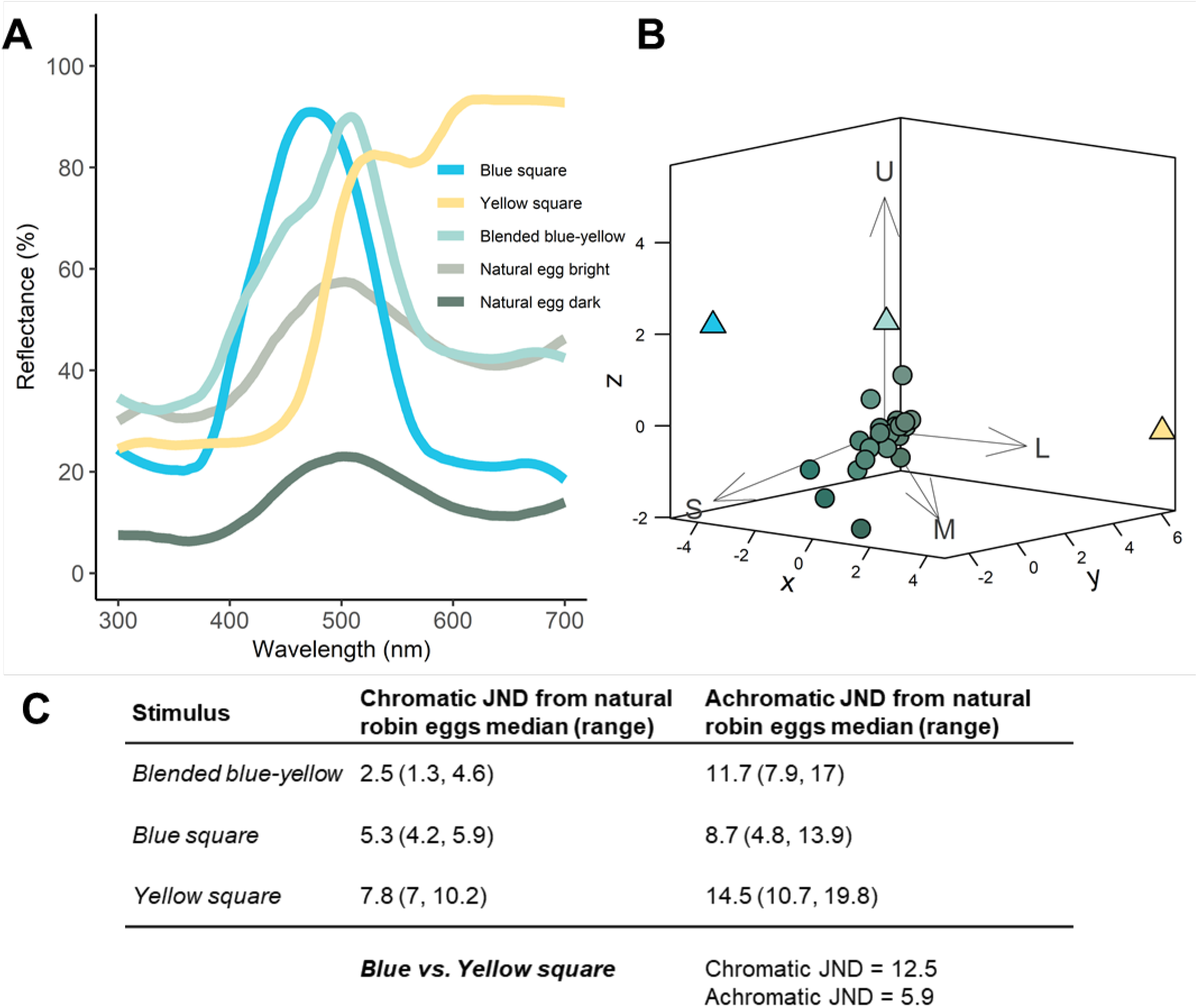
Avian visual modeling of experimental egg colors. **A** Representative reflectance spectra of blue square, yellow square, blended blue-yellow checkered pattern and two natural robin eggs. **B** 3D Euclidian distance Just Noticeable Difference (JND) plot of blue square (triangle), yellow square (triangle), blended blue-yellow checker pattern (triangle) and N=22 natural robin eggs (circles). Relative distance of points to each other are in units of JNDs, arrows represent each avian cone photoreceptor type: *U* = ultraviolet-sensitive cone, *S* = shortwave-sensitive cone, *M* = mediumwave-sensitive cone, *L* = longwave-sensitive cone. Natural robin egg reflectance spectra on the **A** were used from English and Montgomerie (2011) for “bright” and “dark” representative robin egg reflectance spectra, and robin egg reflectance spectra from Croston and Hauber, 2015b were used for **B** and **C** the calculation of JND values.

Previous work has shown that visual modelling of chromatic perceptual differences (i.e., JNDs) using the receptor-noise limited model (Vorobyev and Osorio, 1998) can reliably predict the likelihood of a host recognizing and rejecting a foreign egg (Avilés et al., 2010; Cassey et al., 2008; Honza and Cherry, 2017). Therefore, we chose a “blended” blue/yellow (blue R,G,B = 45, 169, 239; yellow R,G,B = 246, 236, 112) color combination with the lowest median chromatic JND value and range from natural robin eggs (median = 2.51 JND and range = 1.26 to 4.65 JND, Fig. 1). Both achromatic and chromatic JNDs of model egg colors are listed in Figure 1C. Because JND values close to 1 are expected to be more difficult to discriminate between than JND values much greater than 1 (Vorobyev and Osorio, 1998), we predicted robins would perceive our blended blue-yellow model egg as quite similar to natural robin eggs, but not as entirely indistinguishable from them (i.e., imperfectly mimetic). All avian visual modeling was done using the *pavo* v2.3 package (Maia et al., 2019) in *R* v 3.6.1(R Core Team, 2017).

The ability to spatially resolve visual stimuli varies across the avian retina (Fernández-Juricic, 2012). Many bird species, like the American robin, tend to have a center of acuity vision (i.e., fovea) with high spatial resolution ability, surrounded by peripheral retinal area with much lower spatial resolution ability (Moore et al., 2017). In the American robin, the fovea projects into the lateral visual field, whereas portions of the retinal periphery project into the binocular field (PB and EF-J, unpublished data). Peak visual acuity of the robins is estimated to be 14.54 cycles/degree for lateral (foveal) vision and 8.75 cycles/degree for binocular vision (peripheral to the fovea) (using retinal ganglion cell counts and eye axial diameter; PB and EF-J, unpublished data). We used the minimum angle of resolution calculated from robin peak foveal visual acuity 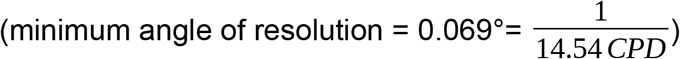 to generate a range of 11 checkered square patterned eggs with visual angles spanning both above and below the minimum angle of resolution at various probable viewing distances (1 to 30cm, Fig. 2A). All visual angles were calculated using Eqn 1.0, where object size is the inter-square distances of the checkered patterns and object distance ranged between 1-30 cm. Viewing distances were roughly estimated as the distance from a robin’s eye, while the robin views an egg using both lateral and binocular vision, to the surface of the model egg using video data of robins viewing eggs from (Hauber et al., 2019) (e.g., see https://youtu.be/GBTML1zcqQA) (Fig. 2A). Checkered pattern designs were all 6 cm × 6 cm in total area and inter-square distances were 0.150 mm, 0.199 mm, 0.299 mm, 0.397 mm, 0.594 mm, 0.845 mm, 0.982 mm, 1.177 mm, 1.463 mm, 1.936 mm, and 2.40 mm. We also created a “control” 0.0 mm pattern using a checkered pattern with 0.150 mm inter-square distances and the Gaussian Blur filter in ImageJ (Schneider et al., 2012) to blend the blue and yellow squares together and create a uniform blue-green colored stimulus (intended to mimic natural robin egg color).

**Figure 2.**
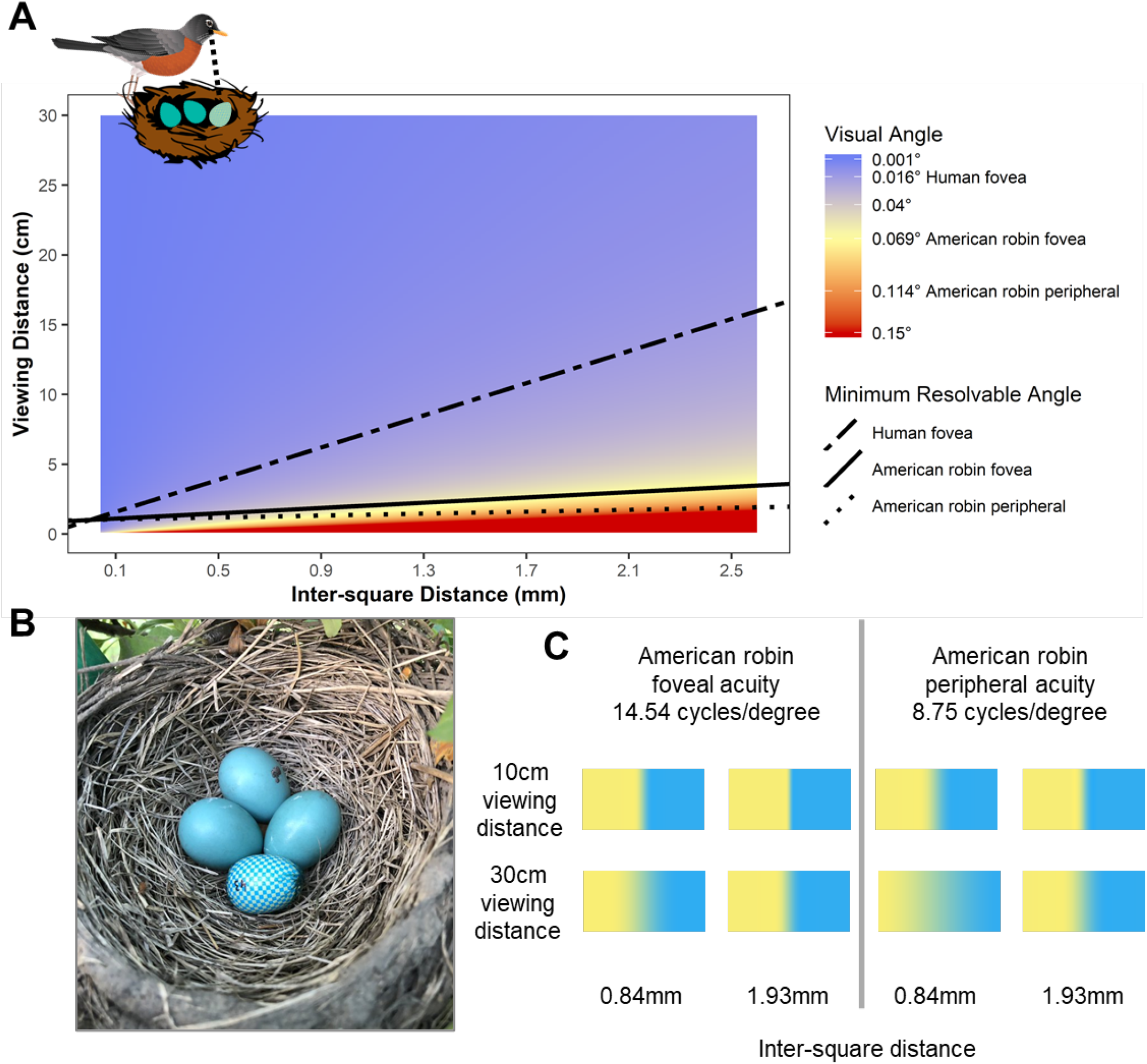
Visual angles of model eggs from the human perspective (dot-dashed line, 60 cycles/degree acuity), and from the American robin’s (*Turdus migratorius*) perspective using either lateral (foveal acuity, 14.54 cycles/degree) or binocular (peripheral acuity, 8.75 cycles/degree) vision. Visual angles were calculated from modelled pattern inter-square distances and viewing distances using *Eqn. 1.0*. We predicted model egg color should appear similar to natural robin egg color when the robin’s visual angle is below the foveal and/or peripheral minimum resolvable angles as visual angles lower than the minimum resolvable angle should cause image blending of blue and yellow squares from the robin’s perspective. Acuity-corrected images were modeled using *AcuityView* image transformations (Caves and Johnsen, 2017) in *ImageJ* (van den Berg et al., 2020).

### Artificial Egg Design

We used 3D printed, brown-headed cowbird-sized model eggs (2.25cm length × 1.69 cm width in size, “Cowbird egg smooth”, purchased from Shapeways, Inc., following Igic et al., 2015) for all experimental model eggs. Cowbird-sized eggs have been used in previous experiments and can be successfully grasped and removed from nests by adult robins (Luro and Hauber, 2017). Experimental egg pattern designs were printed as 6 cm × 6 cm squares onto transferrable decal paper (see *Experimental Egg Pattern* Design above). We cut out, wrapped, and glued the decal patterns onto the model eggs in a vertical orientation towards the egg poles and used small curved scissors to trim away excess decal. Air bubbles were removed by gently pressing the decals by hand until the pattern was flush and smooth on the surface of the model egg. Eggs were left to dry for at least 48 hours before being placed into wild robins’ nests in the field.

### Egg rejection experiments

We searched for active robin nests in Champaign County, Illinois, USA, during May through June of 2019. Upon finding a nest with at least two eggs, we recorded the current clutch size, added a randomly chosen model egg into the nest, and monitored the nest daily until the artificial egg went missing from the nest (rejected) or up to 3 days from egg insertion if the egg remained in the nest (accepted; sensu Luro and Hauber 2017). Adult robins were not captured or marked for this study but we visited three distant sites (more than 5 km apart), used multiple simultaneously active nests within each site, and tested each nest once with a single model egg treatment to reduce biological non-independence. Nest abandonment is not a response to experimental parasitism in robins (Croston and Hauber, 2014), and so data from abandoned (n = 4) or depredated nests (n = 2; out of 33 total nests tested) were not included in the analyses. Sample sizes for each treatment were as follows: control pattern, N = 2; 0.150 mm, N = 2; 0.199 mm, N = 1; 0.299 mm, N = 2; 0.397 mm, N = 3; 0.594 mm, N = 2; 0.845 mm, N = 1; 0.984 mm, N = 2; 1.177 mm, N = 3; 1.463 mm, N = 2; 1.936 mm, N = 2; and 2.40 mm, N = 5.

### Statistical Modelling

We used the *brms* package (Bürkner, 2017) in *R* v3.6.1 (R Core Team, 2017) to run logistic regression models predicting robins’ rejection/acceptance responses to checkered model eggs placed into their nests. Our full model’s predictors included the inter-square distance (mm) of the model egg, the nest clutch size when we inserted the model egg into the nest, and the date at which the experiment was initiated. Our reduced models included the inter-square distance (mm) of the model egg, and either clutch size or date of the experiment. The simplest model included the inter-square distance (mm) of the model egg as the only predictor of robins’ egg rejection responses. We included weakly informative priors (Gelman et al., 2017) for the model intercepts (Student T: df = 3, location = 0, scale = 5) and predictors (Student T: df = 3, location = 0, scale = 5). We ran each model for 10,000 iterations across 4 chains and assessed Markov Chain Monte Carlo (MCMC) convergence using the Gelman-Rubin diagnostic (Rhat) (Gelman et al., 2013). Finally, we used leave-one-out cross validation (LOO) to compare models and determined relative model accuracies for predicting robins’ egg rejection responses using the expected log pointwise predictive density (ELPD) differences between models (Vehtari et al., 2017). The most accurate model is ranked as ELPD = 0 and all other model ELPD values are relative to the best model’s ELPD.

Finally, we determined the approximate inter-square distance rejection threshold (inter-square distance at which rejection probability is 0.5) by simulating posterior fitted values from the most accurate model for experimental egg inter-square distances between 0-2.4mm at 0.001mm increments and calculating the posterior median and 75% credible interval of inter-square distance for posterior fits with rejection probability = 0.5. We used 50% as our arbitrary rejection threshold because the median clutch size for robins nests of our experiments was 3—therefore if robins were only responding to the addition of an egg into their nests and randomly guessing which egg to reject, they would correctly reject our model egg approximately 25% of the time. Thus, using a 0.5 probability of egg rejection as a threshold omits the possibility of random guessing while also selecting for a model egg pattern that is not readily discriminated and rejected (i.e., > 0.5 probability of rejection).

## Results

We used data on the outcomes of the egg rejection experiments at N = 27 different nests with a median of N = 2 experimental replicates per model egg type for all 11 different checkered model egg treatments and the control model egg (0.0 mm inter-square distance). We obtained ≥ 10000 effective samples for each model parameter and all models’ Markov Chains (MCMC) successfully converged (Rhat = 1 for all models’ parameters). American robins’ egg rejection responses were best predicted by a model with egg inter-square distances (mm) as the only predictor (Table 1; ELPD difference = 0). However, there was also fair amount of uncertainty in the model’s predictive reliability (posterior median Bayes *R*^2^ ± median absolute deviation [95% credible interval] = 0.108 ± 0.095 [0.001, 0.272]; where Bayes *R*^2^ is the model predicted variance divided by the sum of model predicted variance and model error variance, see Gelman et al., 2019). Robins’ egg rejection responses increased with larger model egg blue-yellow inter-square distances (mm) (posterior median ± median absolute deviation [95% highest-density interval] = 0.89 ±0.53 [−0.17, 1.97]) (Fig. 3). Finally, we found that the approximate 50% egg rejection threshold for robins’ responses to the blue-yellow checkered model eggs was 1.138mm, corresponding to a visual angle range of 0.114° to 0.0114° from viewing distances of 1 cm and 10 cm respectively, with this approximation having a wide range of credible values (approximate 50% threshold posterior median [75% credible interval] =1.138 mm [0.623 mm, 1.675 mm]). Given a 50% egg rejection threshold pattern inter-square distance of 1.138mm, the pattern should begin to blend from the robin visual perspective using either lateral (foveal) or binocular (peripheral) vision at viewing distances > 3cm.

**Table 1.**
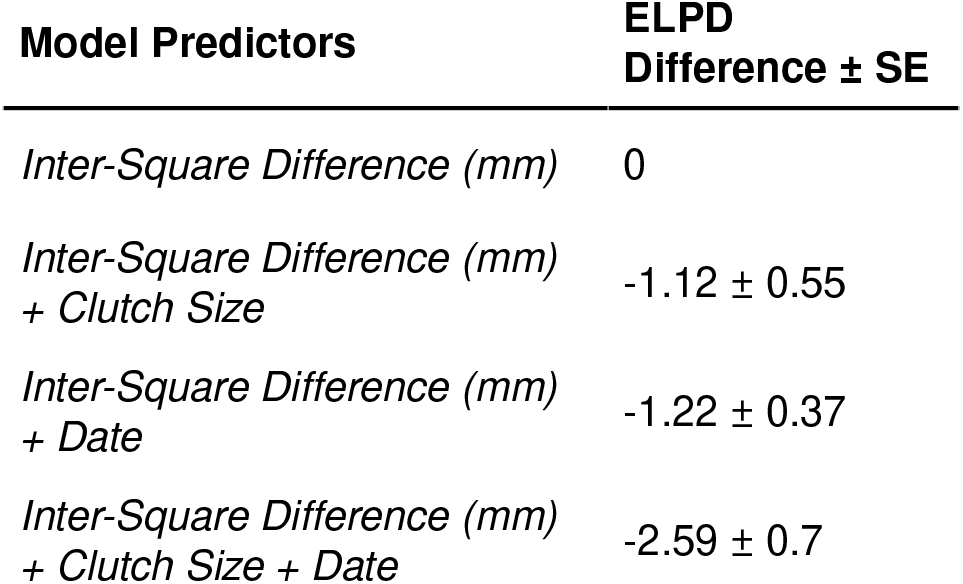
Comparisons of models predicting American robins’ responses to blue-yellow checkered artificial eggs. Models are ordered by their prediction accuracy from leave-one-out cross validation: the top model has an expected log pointwise predictive density (ELPD) difference of 0, and all other models have negative ELPD difference values.

| Model Predictors | ELPD<br>Difference ± SE |
| --- | --- |
| *Inter-Square Difference (mm)* | 0 |
| *Inter-Square Difference (mm)*<br>*+ Clutch Size* | -1.12 ± 0.55 |
| *Inter-Square Difference (mm)*<br>*+ Date* | -1.22 ± 0.37 |
| *Inter-Square Difference (mm)*<br>*+ Clutch Size + Date* | -2.59 ± 0.7 |

**Figure 3.**
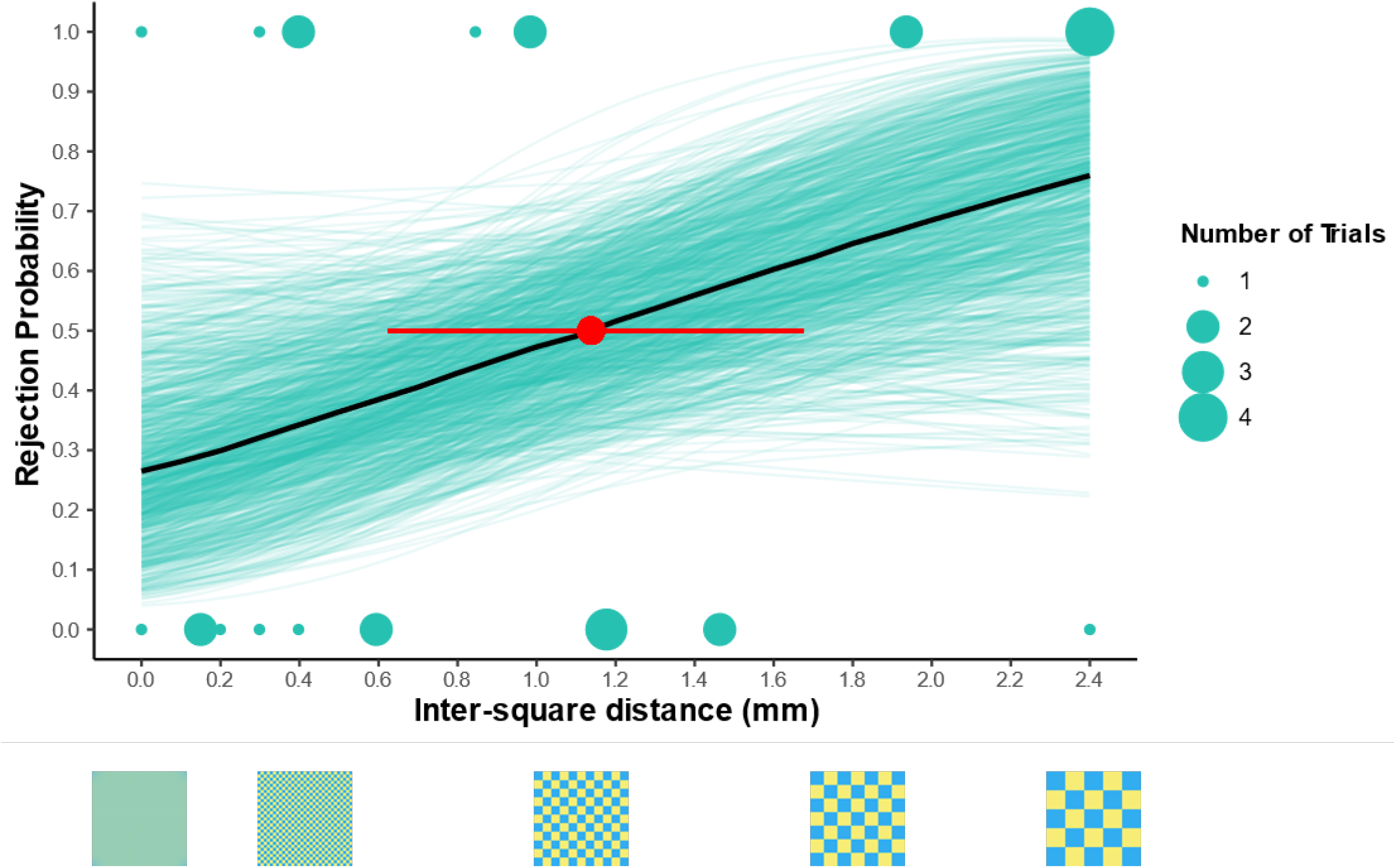
Model-fitted probabilities of American robin egg rejection responses to blue-yellow checkered model eggs. Blue-green lines are simulated model fits for N = 1000 posterior draws; black line is the median of all posterior draws. Model egg inter-square distance between blue and yellow squares at which robins are predicted to reject model eggs at a 0.5 probability is indicated by red 75% credible interval and posterior median. Inter-square distances of several representative checkered pattern images are true to the scale of the x-axis.

## Discussion

Our results suggest that American robins respond predictably to spatial chromatic contrasts when rejecting a foreign egg from the nest. Egg rejection responses were best predicted by inter-square distances of the blue-yellow checkered patterns of model eggs alone (Table 1), as robins were more likely to reject model eggs with larger inter-square distances (Fig. 3). These findings are consistent with the prediction that the combination of blue and yellow colored squares of our model eggs likely appears similar to natural blue-green robin eggs from the robin’s perspective when robins are unable to completely resolve the colored squares as separate features (i.e., when the robin’s visual angle is below the minimum resolvable angle, Fig. 2).

Most passerine birds, including American robins, have laterally placed eyes and often move both their eyes and head when viewing an object of interest to align their high-acuity foveal vision, or align their eyes to gaze at an object using binocular vision with lower acuity perifoveal retinal regions of the eyes (Land, 2014; Moore et al., 2017). The distance at which robins view eggs in the nest, along with their visual acuity when viewing eggs with lateral (foveal) or binocular (peripheral) vision, is equally important for putting our results into context. For instance, if robins consistently inspect their eggs using either lateral or binocular vision from distances > 3 cm, then it is very likely that robins only partially resolved the blue and yellow squares as separate features for all our experimental model eggs (Fig. 2, both foveal and peripheral minimum resolvable angle lines for the American robin). Other factors expected to affect the ability of robins to resolve spatial chromatic contrasts of our model eggs include: pupil size, photoreceptor arrangement and spacing, and potential variation in the spectral profile of light illuminating the eggs and nest (Cronin et al., 2014; Land et al., 2012). Characterizing these visual properties in American robins would make it possible to more precisely test and model robin visual discrimination of egg features. However, visual physiology alone, i.e., modeling visual perception using ocular anatomy and retinal physiology, cannot wholly predict egg recognition and egg rejection behavior of avian brood parasite hosts (Croston and Hauber, 2014; Hanley et al., 2017; Manna et al., 2017; Stoddard and Stevens, 2011). Upon gathering visual information, brood parasite hosts likely use a combination of cognitive decision rules when recognizing and deciding to reject a foreign egg, including counting eggs in the nest (Lyon, 2003), rejecting the most dissimilar egg amongst all eggs in the nest (Moskát et al., 2010), and comparing a foreign egg’s appearance against an internal representation of own eggs’ appearance(Stevens et al., 2013).

Our study also demonstrates the utility of avian egg recognition experiments using model eggs for testing mechanisms fundamental to visually-guided behaviors. Critically, the egg rejection behavior analyzed here is a fitness-relevant behavior for American robins because a small minority of robins do accept brown-headed cowbirds eggs laid in their nests (Lowther, 1981), and raising parasitic cowbird nestlings reduces the fledging success of robin nestlings (Croston and Hauber, 2015a). Surprisingly, many host species that are frequently parasitized by avian brood parasites do not reject foreign eggs laid into their nests even when there is a significant fitness cost to raising unrelated brood parasitic nestlings (Medina and Langmore, 2016). Our results demonstrate that host visual acuity and egg viewing behavior has the potential to limit the ability to detect and discriminate between spatial and chromatic features of eggs. Specifically, hosts can gather more detailed and accurate visual information about egg coloration and patterning by viewing eggs in their nest more often with their high acuity foveal vision and/or viewing eggs from shorter distances to increase the visual angles of the eggs’ patterning. For example, common cuckoos (*Cuculus canorus*) have evolved better egg color and pattern mimicry for host species that have robust egg rejection defenses (Stoddard and Stevens, 2010; Stoddard and Stevens, 2011). Perhaps cuckoo hosts that are skillful egg rejecters have also evolved egg viewing behaviors to better resolve the chromatic and spatial patterning of eggs in their nest, whereas species exhibiting weaker egg rejection responses to foreign eggs have not. Importantly, future experimental studies directly linking predictions derived from visual modelling with behaviors that have known fitness consequences will greatly advance the field of visual ecology (Luro and Hauber, 2020).

Overall, our combined modelling and experimentation revealed that American robin acuity can predict egg rejection responses to foreign eggs, and spatial chromatic contrasts of eggs may be an important visual cue used by birds when viewing and recognizing eggs in their nests. The implication is that visual acuity can impose limits on egg recognition ability in hosts, an important task with considerable evolutionary fitness consequences for hosts of avian brood parasites. Specifically, resolving power constraints of the hosts could be exploited by brood parasites to minimize host detection of parasitic eggs. Finally, failing to recognize differences in visual perception between ourselves and non-human animals is notoriously common and problematic in comparative visual perception studies (Caves et al., 2019). Comprehending differences in visual acuities between ourselves and other species may be especially difficult, considering humans have relatively high visual acuity (Caves et al., 2018). Thus, our results provide a compelling example of how our own biases in the detection, perception and/or processing of visual information may be vastly different from those of birds.

## Ethics Statement

All experimental methods were reviewed and approved by the Institutional Animal care and Use Committee (IACUC) at the University of Illinois (# 17049). Field work was approved by the USA Fish and Wildlife Service (# MB08861A-3) and by the Illinois Department of Natural Resources (NH19.6279).

## Acknowledgements

We thank the residents and landowners of Champaign County for generously allowing us access to American robin nests on their properties.

## Author Contributions

**Alec Luro:** Conceptualization, Methodology, Software, Formal Analysis, Data Curation, Visualization, Writing-Original Draft. **Patrice Baumhardt:** Methodology, Writing-Review & Editing. **Esteban Fernandez-Juricic:** Methodology, Writing-Review & Editing, Supervision. **Mark Hauber:** Conceptualization, Methodology, Investigation, Resources, Supervision, Project administration, Funding acquisition, Writing-Review & Editing.

